# Symmetry breaking transition towards directional locomotion in *Physarum* microplasmodia

**DOI:** 10.1101/675942

**Authors:** Shun Zhang, Juan C. Lasheras, Juan C. del Álamo

## Abstract

True slime mold *Physarum polycephalum* has been widely used as a model organism to study flow-driven amoeboid locomotion as well as the dynamics of its complex mechanochemical self-oscillations. The aim of this work is to quantify the mechanical aspects of symmetry breaking and its transition into directional flow-driven amoeboid locomotion in small (<∼ 200 *µ*m) fragments of *Physarum polycephalum*. To this end, we combined measurements of traction stresses, fragment morphology, and ectoplasmic microrheology with experimental manipulations of cell-substrate adhesion, cortical strength and microplasmodium size. These measurements show that initiation of locomotion is accompanied by the symmetry breaking of traction stresses and the polarization of ectoplasmic mechanical properties, with the rear part of the microplasmodium becoming significantly stiffer after the onset of locomotion. Our experimental data suggests that the initiation of locomotion in *Physarum* could be analogous to an interfacial instability process and that microplasmodial size is a critical parameter governing the instability. Specifically, our results indicate that the instability driving the onset of locomotion is strengthened by substrate adhesiveness and weakened by cortical stiffness. Furthermore, the Fourier spectral analysis of morphology revealed lobe number *n* = 2 as the consistent dominant mode number across various experimental manipulations, suggesting that the instability mechanism driving the onset of *Physarum* locomotion is robust with respect to changes in environmental conditions and microplasmodial properties.

## 1. Introduction

*Physarum polycephalum* is an acellular slime mold long used as model organism of amoeboid motility [1, 2], unconventional computing [3], and biomimetic design of soft robots [4]. In nature, *Physarum* forms branching tubular vein networks up to meters long, which contract rhythmically producing alternating flows of endoplasmic fluid [5]. Sub-millimeter fragments (i.e., microplasmodia) of *Physarum* or droplets of its endoplasmic fluid can be prepared in the laboratory by a variety of methods [6]. Microplasmodia smaller than a few hundred microns do not develop prominent veins and are composed of a submembranous gel-like ectoplasm (i.e., the cortex) layer in addition to the endoplasmic fluid. When seeded onto a substrate, initially round *Physarum* microplasmodia oscillate rhythmically and, in some cases, these fluctuations lead to the elongation of the microplasmodium and its directional locomotion [7]. This transition is generally viewed as a symmetry breaking process [7, 8] but our understanding about the process is still limited. There is data suggesting that only microplasmodia with size larger than ∼ 100 *µ*m can break symmetry [9]. However, the dependence of this critical lengthscale on the mechanical properties of the microplasmodium, or in environmental factors such as substrate adhesiveness, is not well understood. Furthermore, it is unclear how ectoplasm remodeling during symmetry breaking in turn affects the mechanical properties of the microplasmodium or its interactions with the substrate.

Takagi and Ueda experimentally documented the emergence of self-organized non-symmetric cell thickness waves in *Physarum* plasmodia and protoplasm droplets, including standing, traveling and rotational waves [10, 11]. Their work motivated mathematical models of the self-organized waves that considered mechano-chemical feedback between ectoplasm contractions, the endoplasmic flow driven by these contractions, and the transport by the flow of the chemical signal that regulates contractility [12–14]. These models were able to reproduce the experimental observations and predicted two key parameters affecting the emergence the mechano-chemical waves – the drag coefficient between the endoplasm and the ectoplasm, and the Péclet number that quantifies the relative importance of advection versus diffusion in the transport of chemical signals. More recent models have incorporated membrane deformability and substrate adhesiveness to simulate microplasmodial motility during symmetry breaking [15] and directional locomotion [16]. However, these models fail to recapitulate directional locomotion from self-organized mechano-chemical waves – they either produce cell shape oscillations with zero net locomotion [15] or prescribe *ad hoc* traveling wave patterns of active contractility and/or substrate adhesiveness [16, 17].

The partial success of mathematical models of symmetry breaking in *Physarum* microplasmodia implies that crucial aspects of this process are still beyond grasp. This conceptual void is in large part due to a paucity of detailed experimental measurements of the mechanics of symmetry breaking, which has opened a gap between the ability to generate testable ideas by modeling and the ability to experimentally test them, and vice versa. For instance, while existing mathematical models can predict how changing specific parameters (e.g., substrate friction or ectoplasm shear modulus) affects detailed observable variables (e.g., membrane shape or substrate traction stress), the existing experimental data have been until recently mostly limited to measurements of cell thickness over time in round static microplasmodia. Experimental studies of *Physarum* microplasmodial locomotion have become increasingly common in recent years [16, 18–21], but symmetry breaking has comparatively received little attention.

The main objective of the present effort was to shed light into the mechanical aspects of symmetry breaking of *Physarum* microplasmodia, and the subsequent onset of directional locomotion. To this end, we combined measurements of microplasmodium shape and traction stresses with experimental manipulations of substrate adhesiveness, ectoplasm integrity, and microplasmodium size. The experimental data is analyzed to study how a biological system like *Physarum*, break symmetry from its initial stable status and subsequently transit towards directional locomotion, and how this process is affected by these aforementioned mechanical factors. Our measurements show the initiation of locomotion is accompanied by a polarity in cellular mechanical properties and exertion of traction stresses. In addition, our experimental data indicates that instability driving the onset of locomotion is strengthened by substrate adhesiveness and weakened by cortical stiffness. Finally, our result suggests that the initiation of locomotion in *Physarum* is analogous to an instability process, thus provides detailed quantitative data for developing and validating mathematical models.

## 2. Methods

### 2.1. Preparation of Physarum Microplasmodia

*Physarum* polycephalum microplasmodia were prepared on surface culture as previously described [6, 7]. First-generation plasmodia were grown from sporae on 1% agar gel (Granulated; BD) using 150×15–mm culture plates (BD) and fed with oat flakes (QUAKER). The plates were kept in a dark humid environment at room temperature for approximately 1 week. Second-generation fragments approximately 1×1 mm in size were harvested from the growing areas of the first-generation plasmodium and transferred to fresh agar plate, where they were kept overnight. Subsequently, microplasmodia with sizes ranging between ∼ 10*µ*m and ∼ 100*µ*m were excised from the growing tips of the second-generation fragments and placed over collagen coated polyacrylamide (PA) gels embedded with fluorescent microspheres. A 1-mm-thick cap made of a 1% agar gel was placed over the sample with the dual purpose of preventing the PA gel from drying out and gently flattening the microplasmodium to facilitate its visualization, as shown previously [16]. The samples were imaged under the microscope immediately after.

### 2.2. Gel Fabrication

Collagen-coated PA gel pads of 1.5 mm thickness were prepared for traction force microscopy as previously described [22], following established protocols [23]. Each PA gel consisted of a bottom layer without fluorescent beads and a thin (10 µm) top layer containing 1 µm fluorescent beads (FluoSperes; molecular probes) that were used as fiduciary markers to track substrate deformation. The gels were fabricated using 5% acrylamide and 0.3% bisacrylamide (Fisher BioReagents), resulting in a Young’s modulus of 8.73 kPa [24]. The Poisson’s ratio of PA was measured to be 0.46 using elastographic traction force microscopy [25]. PA gels were activated with sulfo-SANPAH (Thermal Scientific) under UV light and coated with 0.15 mg/ml collagen I (Corning).

### 2.3. Microscopy

A Leica DMI 6000B inverted microscope mounted on an automated stage (ASI) and controlled by a PC running Micro-Manager software was used for image acquisition [26]. Three acquisition protocols were followed. Bright-field-only image sequences were recorded over 8 hours under 10X magnification at a 1/30 Hz rate. Traction force microscopy acquisitions were performed both in the bright field and the fluorescent field over 8 hours at a 1/300 Hz rate. Each fluorescent field image consisted of a 40-plane three-dimensional z-stack with an inter-plane spacing of Δ*z* = 1 µm. Particle tracking microrheology acquisitions were performed over 10 seconds under 63X with an oil-immersed lens at 200 Hz rate.

### 2.4. 3D Traction Force Microscopy

The contractile forces generated by *Physarum* microplasmodia induced three-dimensional deformation of their substrate, which was determined by measuring the motion of the fluorescent beads embedded in the PA gel by correlation microscopy. Each instantaneous fluorescent image z-stack was compared with a reference z-stack recorded when the substrate was not deformed, i.e. after the microplasmodia moved out of the field of view. This comparison was carried out by dividing both images into 3D interrogation boxes and maximizing the cross-correlation between each interrogation box and the corresponding interrogation box of the reference image. The maximization was performed using constraints that enforced mechanical equilibrium of forces as previously described [27].

The measured deformation at the top of the substrate was used as boundary condition for the equation of mechanical equilibrium for the PA gel as previously described [22, 28]. Using this solution, we computed the deformation field in the whole polyacrylamide substrate, as well as the traction stress vector 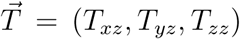 on its top surface. For the sake of presentation, this vector was decomposed into the component parallel to the measurement plane (i.e., in plane), 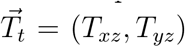, and the out-of-plane component *T*_*n*_ = *T*_*zz*_. The spatial resolution (i.e. the distance between adjacent measurement points) of the traction stress measurements is defined by 1/2-size of the interrogation boxes according to the Nyquist criterion. In the present experiments, the resolution in the *x* and *y* directions was 6.5 µm and 10 µm respectively under 16X and 10X magnification, while it was 20 µm in the *z* direction.

### 2.5. Pharmacological Treatments

In order to stabilize or destabilize the actin filaments of *Physarum*’s ectoplasm, we treated specimens with phalloidin and latrunculin A respectively [29, 30]. Phalloidin treatment was performed by injecting 1 mM of Phalloidin (Sigma) into second-generation molds under a Nikon SMZ-10 microscope using a PM 1000 cell micro-injection system (MicroData Instrument, Inc). The injected amount (100 nl) was less than 1% of the volume of the second-generation mold, as calculated by the measured diameter of the tubular structure of the mold. After waiting for 2 hours, multiple microplasmodia were excised from the marching end as described above. Latrunculin A treatment was performed by transferring microplasmodia to 5 *µ*M Latrunculin A (Cayman) solution for 10 minutes, then washing with Milli-Q water (Millipore).

### 2.6. Modifications of Substrate Adhesiveness

To decrease or increase the substrate adhesiveness we treated the PA gel and agarose cap with Pluronic F-127 or Poly-L-Lysine, respectively [31, 32]. The treatment was performed by soaking the gels in a 0.2% Pluronic F-127 (Sigma) or a 2.5 mg/ml Poly-L-Lysine solution for 1 hour. Subsequently, the gels were washed thoroughly with Milli-Q water three times before seeding the *Physarum* microplasmodia.

### 2.7. Morphological Analysis of Microplasmodia

To quantify the time-dependent shapes of *Physarum* microplasmodia, the perimeter 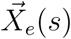 of each microplasmodium was automatically delineated from bright-field images as described previously [28], and its centroid position 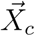 was calculated (see Figure 1A–C). Then the distance from the centroid to each point in the perimeter was measured and expressed as a function of the arclength distance along the contour, i.e. 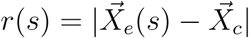 such that *L* = ∮ *ds* is the perimeter length. The resulting function is periodic (i.e. *r*(*s*) = *r*(*s* + *L*)) and its number of peaks provides information about the number of lobes in the microplasmodium perimeter (see Figure 1D–F). Fourier spectral analysis of *r*(*s*) = Σ_*n*_ *R*_*n*_ exp(2*πins/L*) was performed to identify how much each lobe number *n* contributed to departures from circular shape. To this end, we compared |*R*_*n*_|, *n* > 0 with the average radius *R*_0_ = *L*^−1^ ∮ *r*(*s*)*ds*, see Figure 1G–I).

**Figure 1:**
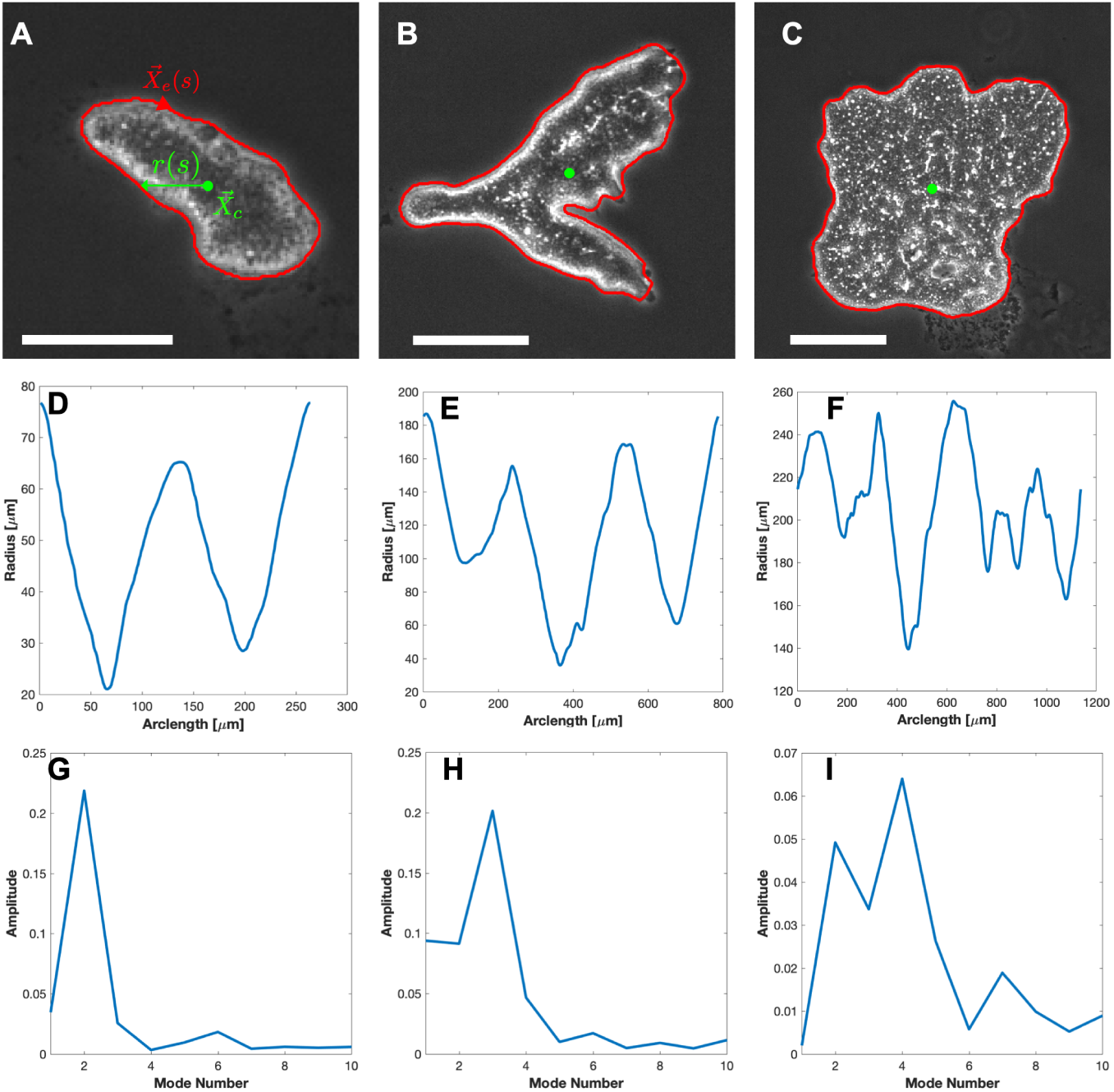
Shape analysis. (**A,B,C**) Snapshots of bright field images of *Physarum* microplasmodia with different morphologies. The red curve is detected outline of the microplasmodium and the green dot is its centroid. (**D,E,F**) Radial distance from the centroid to each point on the microplasmodium’s outline, *r*(*s*), vs. distance along the outline perimeter,*s*. (**G,H,I**) Normalized amplitude |*R*_*n*_|*/R*_0_ of the shape modes derived from the Fourier analysis of *r*(*s*) vs. mode number *n*.

### 2.8. Directional Particle-tracking Microrheology (DPTM)

Carboxylate modified red latex beads with 0.2*µ*m nominal diameter (Fluospheres, Invitrogen, Carlsbad CA) were diluted with Milli-Q water in 1:1000 ratio. The solution was then injected into branched second-generation fragments (as described in Section 2.1) under a Nikon SMZ-10 microscope, using a PM 1000 cell micro-injection system (MicroData Instrument, Inc). The beads were dispersed by shuttle streaming and were evenly distributed across the fragments after 6 hours of injection. At this point, microplasmodia were excised and prepared as described above (Figure 2 A1).

**Figure 2:**
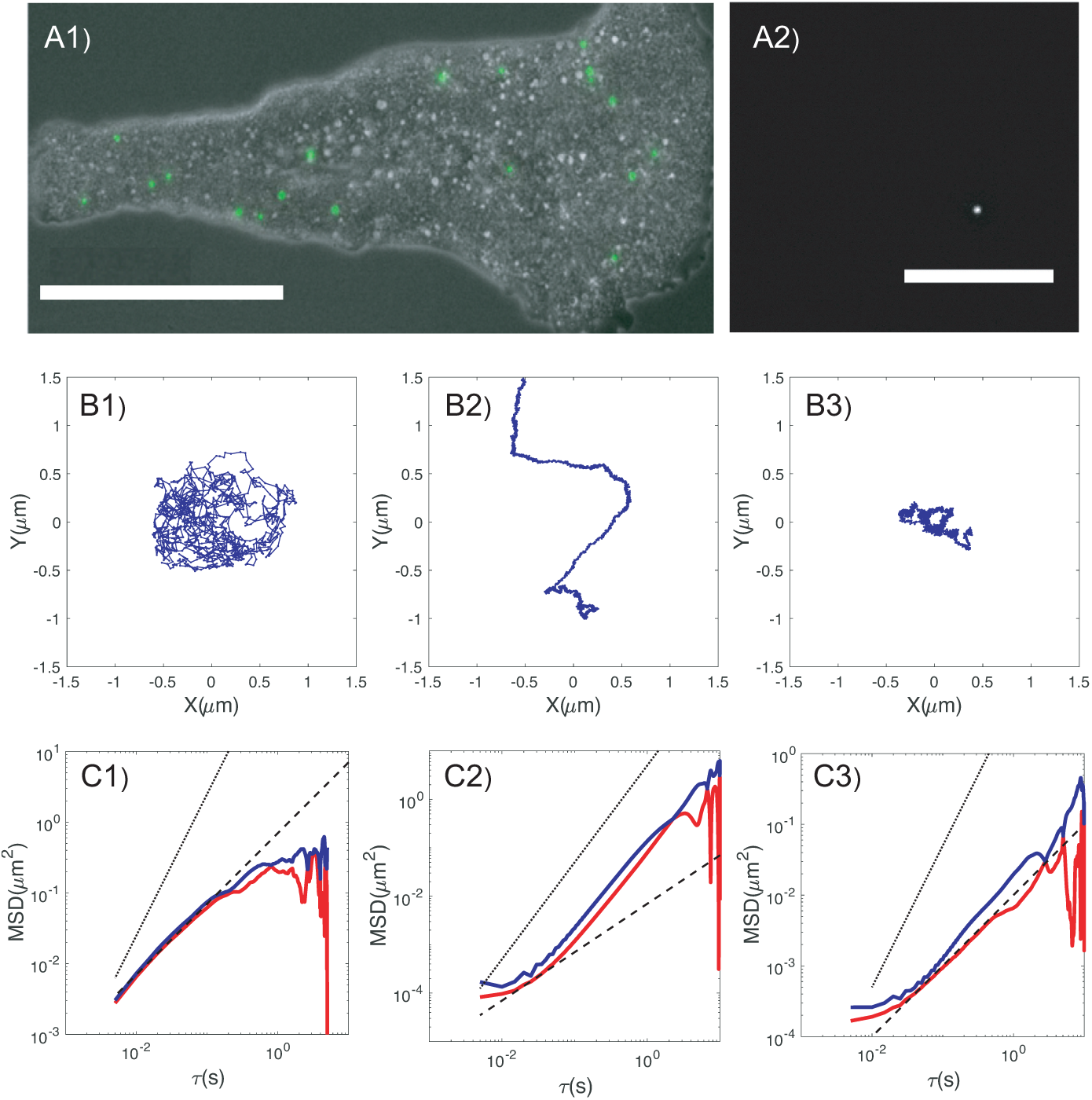
DPTM measurements. (**A1**) Distribution of embedded microparticles in microplasmodial fragment. Scale bar = 100*µ*m. (**A2**) Representative fluorescence image of microparticle acquired in experiment. Scale bar = 20*µ*m. (**B1,B2,B3**) Trajectories of microparticles. (**C1,C2,C3**) Corresponding MSD in the principal directions of maximal (‖) and minimal (⊥) motility for trajectories in (B1–B3). — —, ⟨*x*^2^⟩_‖_; — —, ⟨*x*^2^⟩_⊥_. – – –⟨*x*^2^⟩ ∼ *τ* ….. ⟨*x*^2^⟩ ∼ *τ*^2^;

The motion of the beads embedded in microplasmodia was under the microscope as described above (Figure 2 A2). The centers of the fluorescent beads were tracked using previously described in-house algorithms [33]. The 2 × 2 mean square displacement (MSD) tensor of the beads, ⟨Δ*x*^2^⟩_*i,j*_(*τ*) = ⟨[*x*_*i*_(*t* + *τ*) − *x*_*i*_(*t*)] [*x*_*j*_(*t* + *τ*) − *x*_*j*_(*t*)]⟩ was calculated from their tracked trajectories, and their eigenvalues ⟨Δ*x*^2^⟩_‖_ and (Δ*x*^2^)_⊥_ along the principal directions of maximal and minimal mobility were calculated as previously described [34].

Beads were categorized in three groups according to the shape of their corresponding ⟨Δ*x*^2^⟩_‖_(*τ*) and ⟨Δ*x*^2^⟩_⊥_(*τ*) curves. About 10% of the beads showed isotropic diffusive behavior ⟨Δ*x*^2^⟩_‖_ ≈ ⟨Δ*x*^2^⟩_⊥_ ∼ *τ* at short times that saturated to a constant value *l*^2^ ∼ 1*µm*^2^, suggesting that these beads were trapped in vesicles or membrane invaginations (Figure 2 C1, D1). Another 40% of the beads exhibited ballistic motion with ⟨Δ*x*^2^⟩_‖_ ∼ ⟨Δ*x*^2^⟩_⊥_ ∼ *υ*^2^*τ* ^2^, suggesting that they were being carried by endoplasmic shuttle flow (Figure 2C2, D2). The typical values of *υ* estimated from particle MSDs, *υ* ∼ 0.5*µ* m/s, are in good agreement with previously reported values of endoplasmic velocities in *Physarum* microplasmodia [7, 16]. The remaining ≈ 50% of the tracked beads were diffusive or sub-diffusive in the ⊥ direction (Figure 2C3, D3), suggesting that they were embedded in the F-actin cytoskeleton of the microplasmodium. The ‖ direction of the MSDs for these beads was slightly superdiffusive as a consequence of the slow net directional flow of the ectoplasm, which we previously reported [20]. Thus, we used the measured ⟨Δ*x*^2^⟩_⊥_ for these beads to estimate the shear modulus of the cytoskeleton using the generalized Stokes-Einstein (GSER) relationship [35, 36]. Given that the GSER assumes thermodynamical equilibrium but the MSDs of these beads at timescales longer than *τ* ≈ 0.1*s* are possibly affected by active out-of-equilibrium processes (e.g. contractility of the cytoskeleton) [37, 38], we only considered MSD data with *τ* < 0.1s in our (GSER) analysis.

## 3. Results and Discussion

### 3.1. Plasmodial shape dynamics upon substrate seeding

The size (e.g., average radius *R*_0_, see section 2.7) of *Physarum* plasmodia can be varied across several orders of magnitudes by their method of preparation, resulting in specimens with markedly different morphologies and behaviors. Figure 3 illustrates this dependence in the range 10 *µ*m ≲ *R*_0_ ≲ 1 mm. Upon seeding onto the substrate, *Physarum* fragments with *R*_0_ ≲ 30 µm remained mostly rounded and were unable to initiate directional locomotion despite undergoing considerable shape fluctuations (Figure 3A). Fragments of size *R*_0_ ∼ 100 *µ*m underwent a ∼ 2-hour-long phase of oscillations about a circular shape, followed by a rapid transient in which they adopted a tadpole-like shape and began crawling directionally. Finally, *Physarum* fragments larger than a few hundred microns were also able to crawl directionally but developed a more complex morphology, including a posterior branched vein network (Figure 3C). These fragments are usually called mesoplasmodia as they share some of the key features of both microplasmodia and the larger arborescent plasmodia found in nature. The emergence of their tubular morphology and its effect on locomotion have been previously studied [21, 39], and were beyond the scope of the present study. Here we focus on microplasmodia with homogeneous endoplasmic and ectoplasmic phases (i.e., without tubular networks) in the range of characteristic radius 10 *µ*m ≲ *R*_0_ ≲ 200*µ*m where the transition to locomotion takes place.

**Figure 3:**
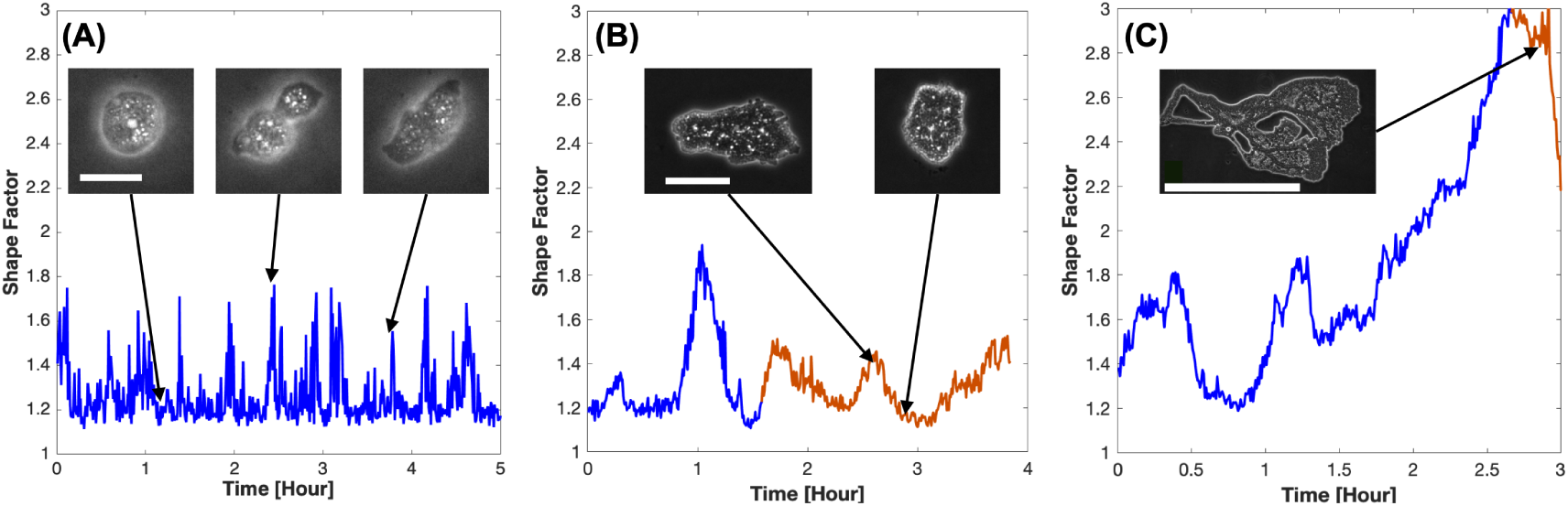
Time histories of the shape factor, 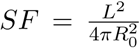, for three *Physarum* plasmodia 0 with different sizes and behavior. Blue color indicates the time interval before the plasmodia initiate locomotion, while orange indicates directional locomotion. The insets show bright field snapshots for the plasmodium corresponding to each *SF* plot. (**A**) Small microplasmodium (scale bar = 20*µ*m) that does not initiate locomotion despite undergoing marked shape fluctuations. (**B**) Larger microplasmodium (scale bar = 100*µ*m) that initiates locomotion approximately 90 minutes after seeding on the substrate. (**C**) Mesoplasmodium (scale bar = 1 mm) that initiates locomotion approximately 150 minutes after seeding on the substrate.

### 3.2. The onset of locomotion is accompanied by a symmetry breaking in traction stresses and ectoplasmic mechanical properties

To better understand the mechanics of the onset of locomotion of *Physarum* microplasmodia, we measured the 3-D traction stresses they exerted on their substrate – both in-plane 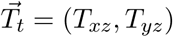 and out-of-plane *T*_*n*_ = *T*_*zz*_. Figure 4 shows a representative example of the time evolution of these stresses during the symmetry breaking process. Initially (*t* ≲ 60*min*), the microplasmodium contracted and relaxed rhythmically while keeping a round shape, similar to the behavior shown in Figure 3. During this phase, it created inward contractile 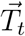 distributed almost uniformly along its body’s contour (Figure 4*a*). At the same time, it exerted intense downward compressive *T*_*n*_ that peaked under the center of the macroplasmodial body (Figure 4*b*). This traction stress pattern is similar to that created by the surface tension and associated Laplace pressure of a vesicle deposited on a soft substrate [40], suggesting that the dynamics of cortical tension could govern the initiation of locomotion in *Physarum*.

**Figure 4:**
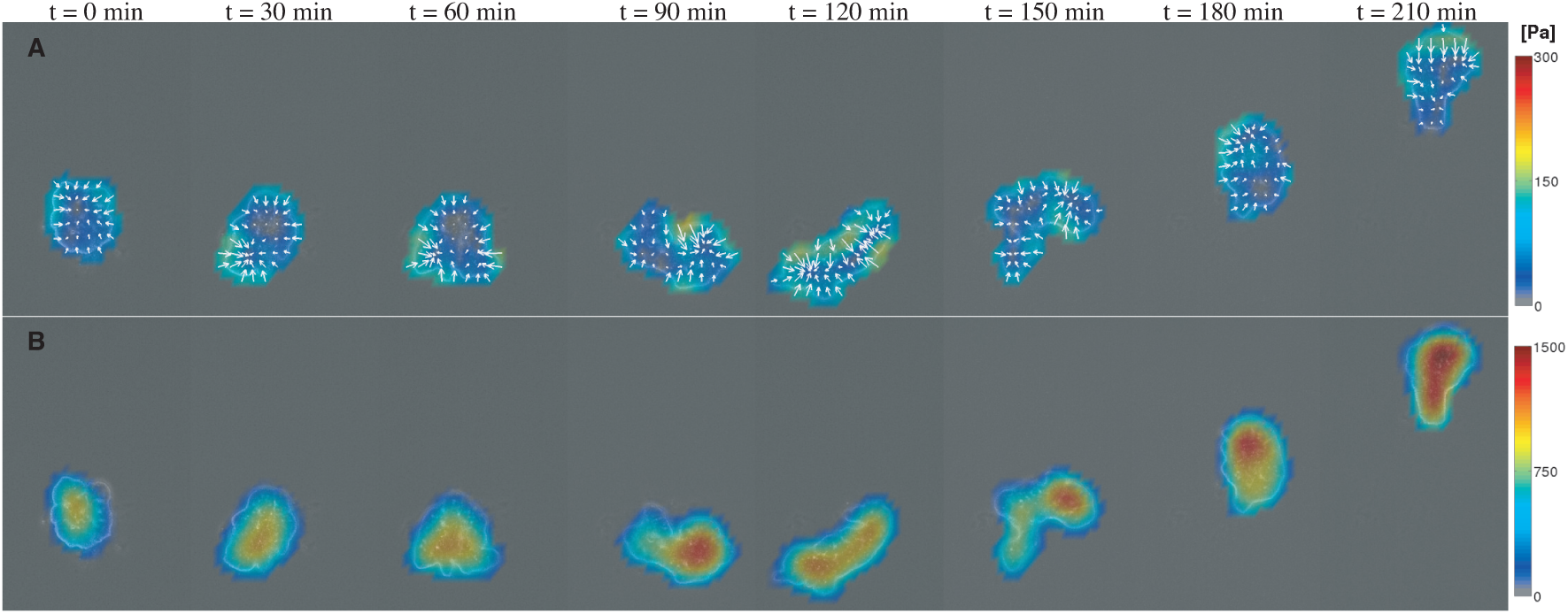
Time evolution of the 3-D traction stresses exerted by a *Physarum* microplasmodium on its substrate during the initiation of locomotion. (*a*) In-plane traction stress vector 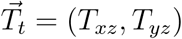; (*b*) Out-of-plane traction stress *T*_*n*_ = *T*_*zz*_.

As the microplasmodium adopted a more elongated shape and started translocating slowly (60 min ≲ *t* ≲ 150 min), the traction stresses lost their initial circular symmetry; however, they had not developed a clear front-back polarity yet. Once the microplasmodium adopted a tadpole-like shape and began directional locomotion (i.e., *t* ≲ 150 min), the traction stresses became more polarized. In particular, the frontal value of *T*_*n*_ became much stronger than its posterior value (Figure 4*b*). This gradient in out-of-plane stresses suggests that the cortex/membrane structure of the microplasmodium bears higher tension in its rear part than in its front. The higher posterior tension would permit a higher pressure difference between the interior and the exterior of the microplasmodium, resulting in lower substrate compression under the plasmodium’s tail. The alternative would be that intracellular pressure was continuously higher at the front of the plasmodium, which would be hard to reconcile with the shuttle streaming flows of alternating directions observed in these microplasmodia [7, 16, 19].

It is possible that the polarization of microplasmodial morphology contributes to polarization in tension, as a tadpole-like shape produces higher membrane curvature at the rear of the microplasmodium. In addition, symmetry breaking may cause long-term polarization in the composition and mechanical properties of the cytoskeleton. The density of myosin II heads in the F-actin cortex is known to increase towards the rear of the microplasmodium [41], and this protein is known to act as a major cross-linker of actin filaments [42]. Moreover, the frontal F-actin cortex undergoes significant remodeling including disassembly and reassembly coinciding with the cyclic contraction and relaxation of the microplasmodium, whereas the posterior cortex has a less dynamic structure [43]. To evaluate the effects of symmetry breaking on the mechanical properties of *Physarum* fragments, we used directional particle tracking microrheology (DPTM) [33–36] to measure the ectoplasmic shear modulus at different locations of the microplasmodium, both before and after the onset of locomotion. In DPTM (see §2.8 for details), the trajectories of beads embedded in the microplasmodium are tracked with high temporal resolution and their 2 × 2 mean squared displacement (MSD) tensor, ⟨Δ*x*^2^⟩_*i,j*_(*τ*) = ⟨ [*x*_*i*_(*t* + *τ*) − *x*_*i*_(*t*)] [*x*_*j*_(*t* + *τ*) − *x*_*j*_(*t*)]⟩, is determined. Based on the shape of the MSDs, we identified particles embedded inside the ectoplasm and discarded particles embedded in vesicles or in the streaming endoplasm. The slow net motion of the ectoplasm was dedrifted by analyzing the smallest eigenvalue of the MSD tensor.

We first measured the MSDs vs. *τ* for ectoplasm-embedded particles at frontal and rear locations of the microplasmodia (Figure 5*a*). For *τ* ≲ 0.1*s*, the MSDs at the rear fell by approximately two-fold, whereas the MSDs at the front did not appreciably change. In contrast, for *τ* ≲ 0.1*s*, the MSDs did not change after symmetry breaking and remained independent of particle position. The *τ*-dependence of these results can be interpreted considering that, for *τ* ≲ 0.1*s*, the random motion of particles embedded inside an actomyosin network is driven by thermal fluctuations so that their MSDs reflect the microrheological properties of the network. On the other hand, for *τ* ≲ 0.1*s* the particle motions reflect network fluctuations driven by motor proteins and the MSDs measure the intensity of this active process [37, 38]. Thus, our results suggest that symmetry breaking causes a change in the mechanical properties of the ectoplasm at the rear part of the microplasmodium, while its contractile activity seems to remain unchanged.

**Figure 5:**
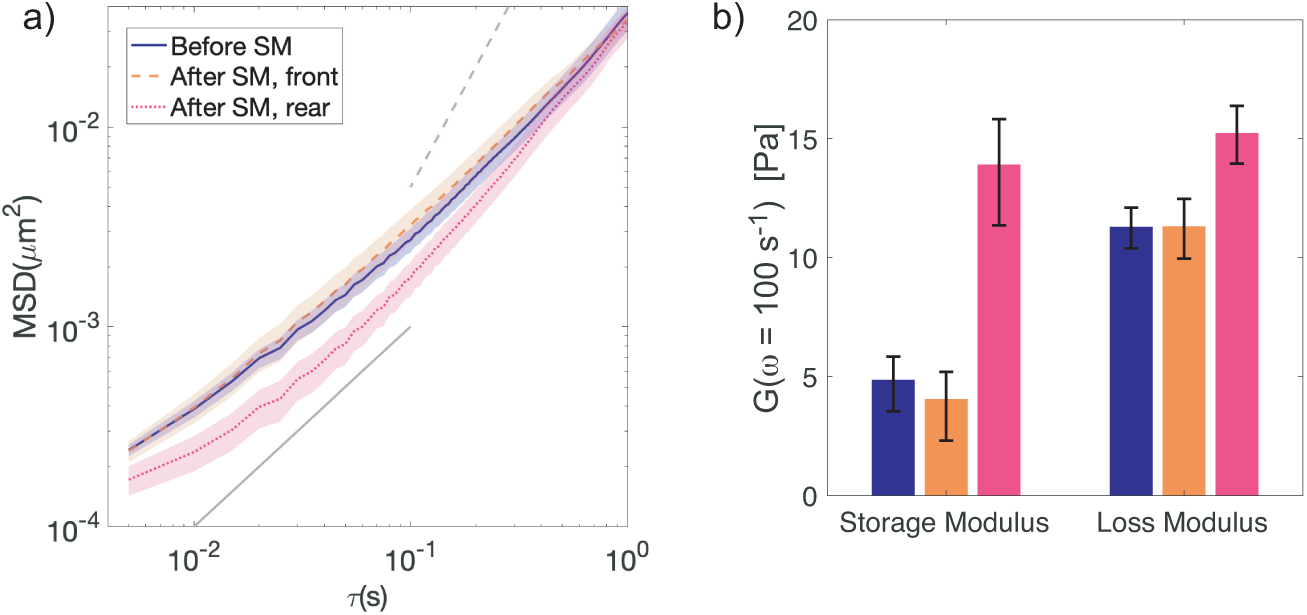
Quantification of the microrheological properties of the ectoplasm in *Physarum* microplasmodia. (*a*) MSD of ectoplasm-embedded particles in the principal direction of minimum mobility (⟨*x*^2^⟩_⊥_). The data come from *Physarum* fragments before initiation of locomotion (N = 42, 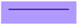), as well as in the head (N = 25,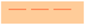) and tail (N = 14, 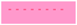) regions of fragments after the onset of directional locomotion. The regions between the average MSD *±* one standard error are shaded in corresponding colors. …….., ⟨*x*^2^⟩_⊥_ ∼ *τ* – – –, ⟨*x*^2^⟩_⊥_ ∼ *τ*^2^; (*b*) Bar plots of storage and loss moduli at 100 Hz calculated from the MSD data in panel (*a*). Error bars correspond to one standard error. The colorscheme is the same as in panel (*a*).

To quantify the changes in ectoplasm microrheological properties after symmetry breaking, we applied the Generalized Stokes-Einstein Relation (GSER, see §2.8 for details) to the measured MSDs in the range 0.005*s* ≤ *τ* ≤ 0.1*s* [35, 36]. This analysis provided loss (viscous) and storage (elastic) shear moduli as a function of frequency *ω, G*″(*ω*) and *G*′(*ω*) respectively. Figure 5(*b*) shows values of *G*′ and *G*″ for *ω* = 100*s*^−1^, indicating that *G*′ at the rear of the microplasmodium increases dramatically after symmetry breaking while *G*″ experiences a moderate increase. Of note, these changes also imply a transition in the material response of the rear ectoplasm from mostly viscous (*G*″ > *G*′) before symmetry breaking to viscoelastic (*G*″ ≈ *G*′) afterward. In contrast, the two components of the shear modulus remained almost constant at the front of the microplasmodium, and their response continued to be viscosity-dominated.

It has long been speculated that the polarization of cortex stiffness may provide a mechanism to select synchronous mechano-chemical wave patterns that facilitate directional locomotion [10, 11]. This idea is based on the observation that protoplasmic droplets with uniform cortical structure often bear rich spatio-temporal mechano-chemical dynamics including chaotic patterns [11], whereas tadpole-shaped locomoting microplasmodia very rarely do so [20]. In fact, prescribing a softer front and a stiffer rear in mathematical models of *Physarum* motility facilitates reproducing experimentally observed behaviors, including directional locomotion [12, 16]. The posterior network of thick-walled veins that develops in mesoplasmodia and larger plasmodia structurally recapitulates a front-back stiffness gradient. Furthermore, it has been shown to act as a low-pass filter that selects organized low-frequency oscillations, facilitating efficient locomotion [21]. As mentioned above, tadpole-shaped microplasmodia also develop a front-back gradient in cortical integrity even if they do not form posterior veins [41]. Our microrheological measurements provide direct evidence that the posterior part of *Physarum* microplasmodia undergo stiffening during the symmetry breaking process that precedes the onset of directional locomotion.

### 3.3. The onset of locomotion is a transition governed by microplasmodium size

To systematically study the role of microplasmodium size in the initiation of locomotion, we tracked the time-dependent shape and position of *N* = 61 microplasmodia of different sizes and categorized them as locomoting or non-locomoting according to their centroid displacement Δ*X*_*c*_ relative to time-averaged radius ⟨*R*_0_⟩_*T*_ over a period *T* = 6 hours, i.e.

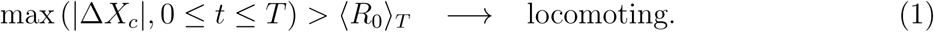

For those microplasmodia that transitioned to directional locomotion, the instant *τ*_*loc*_ at which 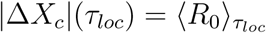 was defined as the time required for initiation of locomotion.

We found that the rate of locomoting microplasmodia increased with ⟨*R*_0_⟩_*T*_ and reached 100 % for *(R*_0_)_*T*_ ≈ 80 *µ*m (Figure 6*a*). This result agrees with Koya and Ueda [9], who studied the emergence of shuttle streaming in microplasmodia formed by coalescence of tiny fragments, each containing approximately eight nuclei, and reported that both the morphological complexity and endoplasmic streaming velocity of the fused microplasmodia experienced a sharp increase beyond a critical size of 20 fragments. Altogether, the available experimental evidence suggests that the initiation of locomotion in *Physarum* is analogous to an instability process and that microplasmodial size is a critical parameter governing the instability. Consistent with this idea, linear stability analyses of mathematical models of *Physarum* as poroelastic mechano-chemical droplets yield growth rates that decrease with decreasing perturbation wavelength [14, 44].

**Figure 6:**
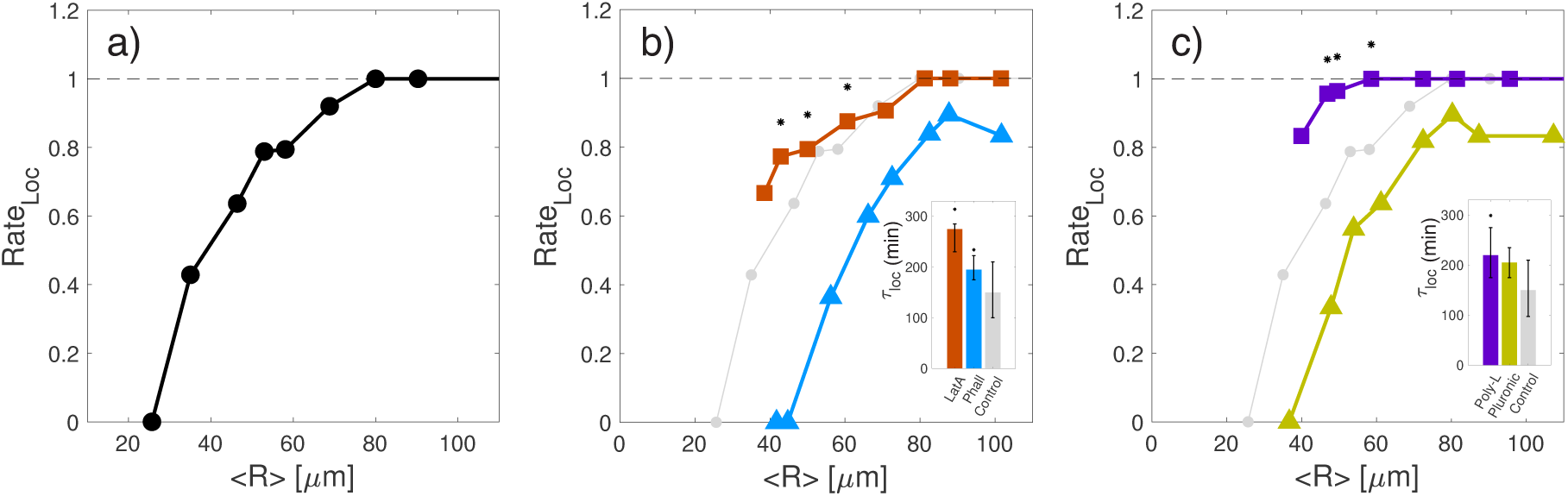
Size, cortical integrity and substrate adhesiveness govern the onset of locomotion. (**a**) Fraction of *Physarum* microplasmodia that initiate directional locomotion during the *T* = 6 hrs ensuing seeding on the substrate, Rate_Loc_, vs. time-averaged microplasmodium size ⟨*R*_0_⟩ (*N* = 61). (**b**) *Rate*_*loc*_ vs. ⟨*R*_0_⟩ for microplasmodia treated with latrunculin A (orange squares, *N* = 70) and phalloidin (blue triangles, *N* = 48). The control condition from panel (*a*) is included with light gray circles for reference. The asterisks denote statistically significant differences between the latrunculin A and phalloidin conditions (*p* < 0.05 according to Fisher’s exact test for categorical variables). (**c**) *Rate*_*loc*_ vs. ⟨*R*_0_⟩ for microplasmodia seeded on substrates coated with poly-L-lysine (purple squares, *N* = 66) and pluronic (yellow triangles, *N* = 39). The control condition from panel (*a*) is included with light gray circles for reference. The asterisks denote statistically significant differences between the poly-L-lysine and pluronic conditions (*p* < 0.05 according to Fisher’s exact test for categorical variables). The inset barplots in **b–c** indicate the median time for initiation of locomotion in each case (error bars denote 95% confidence intervals), *τ*_*loc*_. The asterisks denote statistically significant differences with respect to the control case (*p* < 0.05 according to Wilcoxon ranksum test).

### 3.4. The onset of directional locomotion is governed by ectoplasm integrity and substrate adhesiveness

Next, we studied how the onset of locomotion is affected by the mechanical properties of the microplasmodium and its environment. We first investigated the effect of ectoplasm integrity by pharmacological manipulation of the F-actin cytoskeleton. *Physarum* has a thick cortical ectoplasm composed by F-actin that surrounds the interior of its membrane and is crucial for the generation of the forces and shape changes required for locomotion [45]. To strengthen the ectoplasm, we treated microplasmodia with Phalloidin, which binds to actin filaments and stabilizes them by inhibiting their depolymerization, thereby promoting cortical thickening [46]. To disrupt the ectoplasm, we treated microplasmodia with Latrunculin A (LatA), which prevents actin monomers from polymerizing and has been shown to weaken the cortex in *Physarum* [47]. For each type of treatment, we tracked a large number of microplasmodia for *T* = 6 hours and determined the rate of locomoting specimens as a function of their size, similar to Figure 6(*a*). These experiments revealed that phalloidin-treated microplasmodia of a given size are less prone to initiating locomotion than LatA-treated and control microplasmodia of the same size (Figure 6*b*). Similar to the control case, Phalloidin-treated microplasmodia transitioned from non-locomoting to locomoting as their size increased, but this transition shifted by ≈ 20 *µ*m.

If one idealizes the microplasmodia as a two-phase active poroelastic material, the observed behavior could be explained by the fact that for the same level of contractile force, a stiffer gel phase (ectoplasm) would tend to deform less and squeeze less sol phase (endoplasm). Consistent with this idea and our experimental results, the most unstable wavelength in linear stability analyses of *Physarum* droplets increases with gel phase stiffness [44]. One could also consider that the F-actin network in Phalloidin-treated microplasmodia likely has a smaller pore size, which would cause an increase in the drag coefficient between the gel and sol phases. This effect would damp the growth rate of perturbations, but it would also decrease the wavelength of the most unstable perturbations [14], which is at odds with our experimental data showing that the critical size for symmetry breaking increased with phalloidin treatment (Figure 6*b*).

An alternate explanation for our experimental observations could be that the interfacial tension and bending stiffness of the cortex/membrane system hinder the initiation of locomotion by damping shape perturbations that increase local cortex curvature (e.g., pseudopod protrusions). In consonance with Figure 6(*a*)–(*b*), this mechanism would make microplasmodia of smaller size less likely to initiate directional locomotion. Also in consonance with that figure, this mechanism would make microplasmodia with stronger (weaker) cortices – i.e., Phalloidin-treated (LatA-treated) – more (less) stable than controls. Of note, in smaller *Dictyostelium* amoeboid cells, genetic manipulation of cortex-specific proteins has shown that strengthening cortical strength limits cell migration speed [40]. We are not aware of mathematical models that specifically investigate the effect of cortical integrity in the initiation of *Physarum* migration. However, researchers have explored the general idea of interfacial instabilities in single-phase active fluids [48, 49], offering insight that is at least qualitatively applicable to our experiments. These studies suggest that initially-round cell fragments can become unstable if their size exceeds a critical radius *R*_*crit*_ and that cortical tension and bending stiffness can shift the instability to larger radii. Both predictions are in qualitative agreement with our experimental data (Figure 6*a*–*b*).

An additional qualitative prediction of interfacial instability models is that increasing substrate adhesiveness should contribute to destabilizing microplasmodial shape [48] – similar to increasing the capillary number of the interface of a passive droplet [50]. To test this hypothesis, we experimentally modulated the effect of substrate adhesiveness by performing experiments on substrates coated with either Pluronic F-127 or Poly-L-Lysine. Pluronic F-127 is a bio-compatible PEG-based triblock copolymer surfactant that reduces specific and non-specific adhesion [31]. Poly-L-Lysine is a positively charged polymer that increases substrate adhesiveness by electrostatic interactions with the negatively charged cell membrane [32, 51]. Our experimental results (Figure 6*c*) indicate that microplasmodia of similar size are more likely to initiate locomotion on more adhesive substrates, whereas the opposite is true for less adhesive substrates. Specifically, the Rate_*Loc*_ curves in Figure 6(*c*) are shifted in ⟨*R*_0_⟩ by approximately −20 *µ*m and +15 *µ*m for Poly-L-Lysine and Pluronic F-127 treated substrates, respectively.

While the measured Rate_*Loc*_ curves are in good qualitative agreement with previous model predictions, the measured times for initiation of locomotion (*τ*_*loc*_, see insets in Figure 6*b*-*c*) call for caution when interpreting our experiments in terms of linear stability of small perturbations. The data indicate that, as expected, those experimental manipulations that stabilized microplasmodia around their initially round shape (i.e., Phalloidin treatment and Pluronic substrate coating) led to an increase in *τ*_*loc*_. However, the manipulations that destabilized microplasmodia (i.e., LatA treatment and Poly-L-lysine substrate coating) led to even more substantial increases in *τ*_*loc*_, which is in apparent contradiction with the faster perturbation growth that one would expect in those conditions. This discrepancy brings up the limitations of linear stability analysis – predicting the long-term growth of shape perturbations usually requires full-blown simulations incorporating non-linear interfacial mechanics, substrate interactions, and mechano-chemical feedback. But it also highlights that experimental manipulations of biological systems usually have downstream effects on multiple parameters. For instance, weakening the F-actin cytoskeleton can cause a decrease in actomyosin contractile forces that drive cell shape changes [52]. Besides, increasing substrate adhesiveness can increase membrane tension by increasing the area of contact between the plasmodium and the substrate, which causes membrane stretching [53].

To evaluate the effect of our experimental manipulations in *Physarum* microplasmodia, we measured the average magnitudes of the in-plane and out-of-plane traction stresses (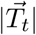 and |*T*_*n*_| respectively) for LatA, phalloidin, Poly-L-Lysine and pluronic treatments, and for control conditions. We also determined the correlation between microplasmodial spread area and in-plane traction stress magnitude, 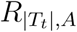 under these different conditions. The latter is an interesting parameter because it provides information about the role of mechanical forces in driving microplasmodial shape oscillations. In our experiments, LatA treatment did not lead to significant changes in 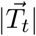 but it caused a moderate decrease in |*T*_*n*_| compared to control, consistent with reduced contractility of a slightly weaker actomyosin cytoskeleton (Figure 7*a*-*b*). This reduced contractility probably explains why *τ*_*loc*_ increased in LatA treated microplasmodia (Figure 6*b*, inset). Phalloidin treatment caused strong increases in both 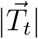 and |*T*_*n*_|. While this increase is compatible with the intended manipulation in cortical integrity, the resulting higher contractility may explain why *τ*_*loc*_ only grew moderately with this manipulation (Figure 6*b*, inset). Substrate coating with pluronic did not significant affect |*T*_*n*_| or 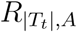 (Figure 7*b, c*) but it significantly decreased |*T*_*t*_| (Figure 7*a*), which is to be expected given that this treatment decreases substrate friction. Finally, substrate coating with poly-L-lysine significantly increased |*T*_*t*_| without significantly increasing |*T*_*n*_| (Figure 7*a, b*), consistent with the intended effect of the manipulation – to increase substrate friction. However, this coating also resulted in an almost total loss of correlation between |*T*_*t*_| and spread area (Figure 7*c*), implying that the increased friction made it harder for the microplasmodium to change its shape by means of contractile stresses. This result may explain the dramatic increase in τ_*loc*_ observed for poly -L-lysine (Figure 6*c*, inset), even if this manipulation overall favored symmetry breaking at smaller microplasmodial radii.

**Figure 7:**
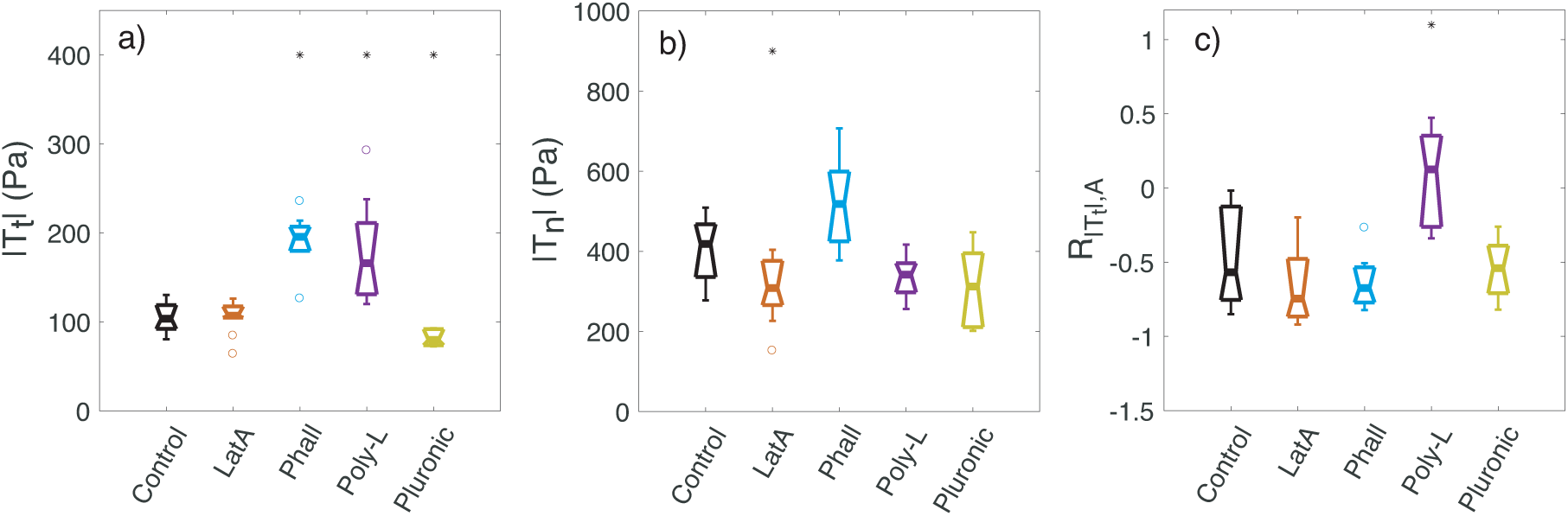
Cortical integrity and substrate adhesiveness affect the exertion of traction stresses. (**a**) Box plots of the mean magnitude of in-plane traction stresses 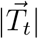 exerted by microplasmodia treated with latrunculin A (*N* = 13) or phalloidin (*N* = 8), or seeded on substrates coated with poly-L-lysine (N=9) or pluronic (N=5), together with untreated microplasmodia seeded on collagen coated substrates (control, *N* = 8). (**b**) Box plots of the mean out-of-plane traction stresses magnitude |*T*_*n*_| for the same microplasmodia in panel (*a*). (**c**) Box plots of Spearman correlation coefficient between 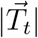 and and microplasmodial spread area. The asterisks denote statistically significant differences with respect to control conditions (*p* < 0.05 according to Wilcoxon’s ranksum test).

### 3.5. Dominant Shape Oscillation Modes

As noted above, *Physarum* microplasmodia of all sizes underwent pronounced shape fluctuations upon seeding onto the substrate (see Figure 3). Figure 8(*a*) shows the distributions of shape modes as a function of lobe number obtained from the morphological analysis described in section 2.7, after normalizing with *R*_0_ and averaging over time. The data indicate that the most dominant lobe number in *Physarum* microplasmodia that initiated locomotion was *n* = 2, which in combination with the prominent albeit less dominant mode *n* = 1 corresponds to an oval/tadpole-like shape (see Figure 1*a*). The second most dominant mode was *n* = 3, which in combination with modes *n* = 2 and *n* = 1 yields a tadpole-like shape with two frontal pseudopods (see Figure 1*b*). Remarkably, lobe number *n* = 2 was consistently dominant for 98% of the microplasmodia (281 out of 287) both in control conditions and in experiments where we manipulated the F-actin cytoskeleton and substrate adhesiveness. Together with the similar shapes of the *Rate*_*Loc*_ vs. *R*_0_ curves observed in Figure 6 for all conditions, this result suggests that all the microplasmodia in our experiments underwent a similar instability mechanism to initiate locomotion.

**Figure 8:**
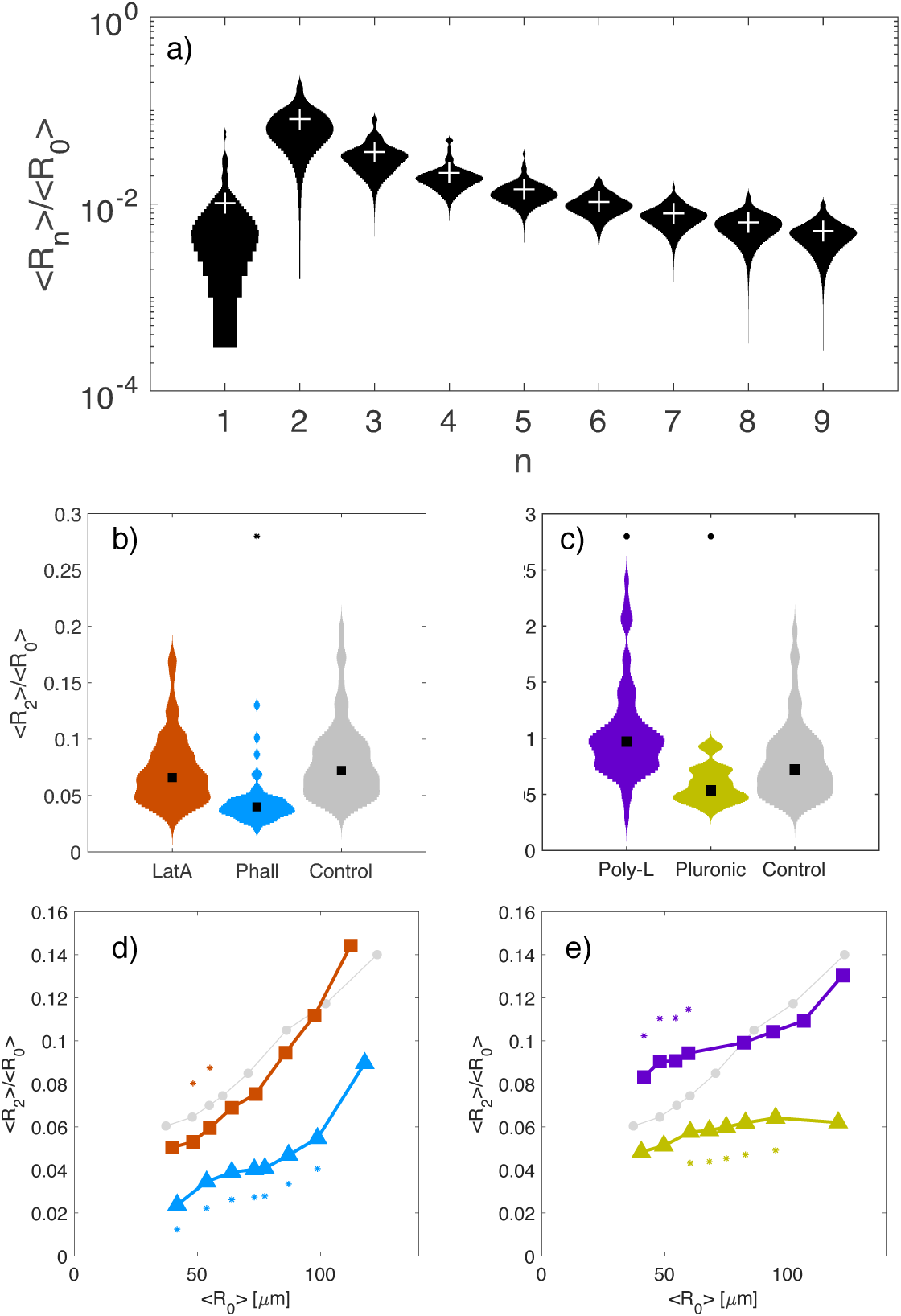
Size, cortical integrity and substrate adhesiveness determine the morphology of fragments. (**a**) Distributions of normalized shape mode amplitudes ⟨*R*_*n*_⟩*/* ⟨*R*_0_⟩ vs. lobe number. Normalized amplitude of the dominant shape mode, ⟨*R*_2_⟩*/* ⟨*R*_0_⟩ for microplasmodia treated with latrunculin A (*N* = 70), phalloidin (*N* = 48) as well as untreated ones (*N* = 61). ⟨*R*_2_⟩*/* ⟨*R*_0_⟩ for microplasmodia seeded on substrates coated with poly-L-lysine (*N* = 66), pluronic (*N* = 39) and collagen (control, *N* = 61). (**d**) ⟨*R*_2_⟩*/* ⟨*R*_0_⟩ vs. ⟨*R*⟩ for the three same cases as panel (*b*). (**e**) ⟨*R*_2_⟩*/* ⟨*R*_0_⟩ vs. ⟨*R*⟩ for the three same cases as panel (*c*). The asterisks denote statistically significant differences with respect to control conditions (*p* < 0.05 according to Wilcoxon’s ranksum test).

Figure 8(*b*)-(*c*) shows that the dominant shape perturbations became significantly weaker in Phalloidin-treated microplasmodia and in substrates coated with Pluronic, whereas they became significantly stronger in Poly-L-lysine coated substrates. These results provide additional evidence suggesting that the instability driving the onset of locomotion is strengthened by substrate adhesiveness and weakened by cortical stiffness. Furthermore, Figure 8(*d*)-(*e*) indicates that the amplitude of the dominant shape perturbations increased with ⟨*R*_0_⟩, particularly beyond values approximately equal to the critical radius of initiation of locomotion observed in Figure 6. Microplasmodia seeded onto Pluronic-treated substrates deviated from this behavior, suggesting that larger size may not confer an important advantage for locomotion in less adhesive substrates.

## 4. Conclusion

The plasmodium *Physarum* polycephalum, a true slime mold, has been widely used as a model organism in studies of flow-driven amoeboid locomotion [1,2,12]. The plasmodium is a single cell with multiple nuclei and is composed of a gel-like submembranous ectoplasmic layer and a sol-like inner endoplasm. The endoplasm exhibits streaming flows of alternating direction induced by oscillatory contractions of the ectoplasm, which are in turn governed by the flow-mediated transport of chemical signals such as calcium ions [7,16,20]. A remarkable feature of this kind of flow-contraction coupling is that it generates long-range self-organization [54]. Consequently, the size of *Physarum* plasmodia can vary over several orders of magnitude without drastic qualitative changes in their overall behavior [6, 9]. In contrast, their locomotor dynamics can be rather sensitive to plasmodial size. Particularly, *Physarum* microplasmodia of sizes smaller than ∼ 80 microns do not migrate directionally whereas microplasmodia of a few hundred microns undergo rapid amoeboid locomotion [7, 9, 16].

The transition between the non-migratory and the migratory states of *Physarum* microplasmodia is reminiscent of a symmetry breaking process [7,8]. Given the biological simplicity of *Physarum* and its very loose control over specimen size, this organism constitutes an excellent model to study symmetry breaking phenomena in biological active matter. More specifically, experiments with small *Physarum* microplasmodia of sizes ranging from tens to a few hundred microns allow for studying flow-driven amoeboid motility without the posterior vein structures usually found in fragments of larger size (e.g., mesoplasmodia [21]). However, the mechanics of symmetry breaking and the onset of locomotion in *Physarum* microplasmodia had not been studied experimentally before.

This study has revealed that substrate adhesiveness, microplasmodial size, and cortical strength govern the symmetry breaking and onset of locomotion of *Physarum* microplasmodia. Overall, our results are consistent with two not mutually exclusive views of the initiation of locomotion – one that idealizes the process as the instability of a two-phase active poroelastic material [14, 15, 44], and an alternate view that models symmetry breaking as an interfacial instability [48, 49]. In both views, the integrity of the ectoplasm hinders the initiation of locomotion at short length scales, although by different physical mechanisms. In both scenarios, increased substrate friction favors the onset of locomotion of small fragments that would otherwise remain round and stable, consistent with our experiments on pluronic- and poly-L-lysine-coated substrates. However, we note that increased substrate friction also caused an increase in the experimentally observed timescale of symmetry breaking, suggesting that indefinitely increasing substrate adhesion should arrest *Physarum* locomotion, consistent with our previous observations [16].

Our spectral analysis of microplasmodial shape during symmetry breaking revealed that the onset of locomotion coincides with a transition from a round shape to a tadpole-like shape with two lobes or, albeit less frequently, three lobes. This result was not sensitive to manipulations in ectoplasm integrity or substrate adhesiveness, suggesting that the underlying mechano-chemical instability was the same for different microplasmodial sizes and experimental manipulations. Our measurements also revealed that an intense polarization of the mechanical properties of the ectoplasm accompanied the onset of locomotion – over ∼ 3 hours, the elastic shear modulus of the posterior ectoplasm increased approximately three-fold, whereas its viscous modulus increased only by ∼ 50 %. The robustness of the mechano-chemical instability triggering directional locomotion and the anteroposterior polarization of ectoplasmic mechanical properties may facilitate the emergence of mechano-chemical wave patterns leading to directional locomotion, as previously speculated by [10, 11]. Consistent with these ideas, disorganized mechano-chemical waves are a seldom occurrence in directionally migrating *Physarum* microplasmodia [20]. However, the present data could not clarify the previously reported emergence of different organized wave patterns upon symmetry breaking – i.e., traveling waves (*peristaltic* locomotion) and standing waves (*amphistaltic* locomotion) [7, 16, 19, 20]. Given that *peristaltic* locomotion is significantly faster than *amphistaltic* locomotion, this question warrants future investigation.

In summary, the present study suggests that a mechano-chemical instability governs the onset of locomotion in *Physarum* microplasmodia. The nature of the instability does not seem to depend on substrate adhesiveness, microplasmodial size, or cortex integrity; however, the observed growth rates are sensitive to these factors. Specifically, the initiation of locomotion takes place when microplasmodial size is larger than a critical value, and this critical size decreases with increasing substrate adhesiveness or weakening cortex integrity. The posterior cortex becomes significantly stiffer concurrent to this process, which presumably favors directional locomotion by preventing the growth of shape fluctuations at the tail of the microplasmodium. While these experimental measurements leave open questions about how mechanical factors work together to enable flow-driven amoeboid locomotion, they also provide original quantitative data with an unprecedented level of detail that can be used to inspire and validate future modeling studies aimed at addressing those questions.

